# Pseudogenes as a neutral reference for detecting selection in prokaryotic pangenomes

**DOI:** 10.1101/2023.05.17.541134

**Authors:** Gavin M. Douglas, B. Jesse Shapiro

## Abstract

A long-standing question is to what degree genetic drift and selection drive the divergence in rare accessory gene content between closely related bacteria. Rare genes, including singletons, make up a large proportion of pangenomes (the set of all genes in a set of genomes), but it remains unclear how many such genes are adaptive, deleterious, or neutral to their host genome. Estimates of species’ effective population sizes (N_e_) are positively associated with pangenome size and fluidity, which has independently been interpreted as evidence for both neutral and adaptive pangenome models. We hypothesised that pseudogenes, used as a neutral reference, could be used to distinguish these models. We find that most functional categories are depleted for rare pseudogenes when a genome encodes only a single intact copy of a gene family. In contrast, transposons are enriched in pseudogenes, suggesting they are mostly neutral or deleterious to the host genome. Thus, even if individual rare accessory genes vary in their effects on host fitness, we can confidently reject a model of entirely neutral or deleterious rare genes. We also define the ratio of singleton intact genes to singleton pseudogenes (s_i_/s_p_) within a pangenome, compare this measure across 668 prokaryotic species, and detect a signal consistent with the adaptive value of many rare accessory genes. Taken together, our work demonstrates that comparing to pseudogenes can improve inferences of the evolutionary forces driving pangenome variation.

## Introduction

Bacterial strains within the same species often encode substantially different genes. This has been established through genome analyses where the entire set of ubiquitous (‘core’) and variably present (‘accessory’) genes across strains are taken to encompass a single ‘pangenome’. Based on such analyses, the percentage of accessory genes within pangenomes varies from 40-80% across prokaryotic species^1^. Many individual accessory genes have been shown to be adaptive, but it remains controversial whether genetic drift or natural selection are responsible for driving overall pangenome variation across species.

This has been investigated by comparing species pangenome diversity to measures of effective population size (N_e_). N_e_ represents the population size under idealized conditions and is the key parameter determining the efficacy of selection vs. genetic drift. Several methods can be used to estimate, or be used as proxies for, N_e_. One common proxy is the ratio of non-synonymous to synonymous substitution rates (dN/dS) in core genes. Assuming stronger purifying selection against non-synonymous mutations relative to synonymous mutations in core genes, a lower dN/dS ratio indicates higher selection efficacy and thus a higher N_e_. Lower dN/dS ratios are associated with higher pangenome diversity across prokaryotic species^2,3^, which has been interpreted as evidence for higher selection efficacy to retain slightly adaptive accessory genes, resulting in larger pangenomes^3,4^. If there was strong selection acting on most accessory genes, then they would be retained regardless of species N_e_ (except perhaps for those with very low N_e_).

One issue with this adaptive explanation is that neutral genetic variation is also higher in species with higher N_e_ due to weaker genetic drift^5^ (e.g., fewer population bottlenecks). Indeed, nucleotide diversity at neutral sites is the standard method for calculating N_e_, assuming equal mutation rates across species. Pangenome diversity is also positively associated with N_e_ based on this approach^6^. Accordingly, neutral and adaptive explanations for pangenome diversity cannot be distinguished based on associations with N_e_ alone, especially for rare accessory genes that could represent segregating neutral variation. Others have remarked that this remains an issue as it is unclear how to partition genes into categories expected to experience distinct selective pressures^7^.

We hypothesised that pseudogenes – genes degenerating through the introduction of mutations such as premature stop codons, insertions, and deletions – could be used as an approximately neutral reference for detecting selection on rare intact accessory genes. Pseudogenes can arise when genetic drift overcomes purifying selection to retain a gene^8^, or through positive selection to eliminate a deleterious gene^9^. We reasoned that accessory gene families that tend to remain intact are likely under stronger purifying selection than those that tend to be pseudogenized. This insight is particularly relevant for rare accessory genes, which make up the largest fraction of pangenomes^10^, and for which the evolutionary forces are most controversial. We investigated this idea by first comparing the functional categories of rare intact genes and pseudogenes across ten well-sampled bacterial species. We then conducted a broader assessment of intact genes and pseudogenes in the pangenomes of nearly 700 different prokaryotic species.

## Results

### Comparing functional annotations of rare genes and pseudogenes across well-sampled species

If rare accessory genes were effectively neutral to host fitness, then we would expect no difference in the functional annotations between intact genes and pseudogenes within a species. In contrast, differences in functional annotations between these rare element types could suggest functions that tend to be beneficial in a genome.

We conducted a comparison of the functional annotations of rare intact genes and pseudogenes in a dataset of 10 bacteria species with a relatively high number of sequenced genomes (135-6,845 genomes per species), including highly sampled human pathogens and bacteria with other lifestyles: *Agrobacterium tumefaciens, Enterococcus faecalis*, *Escherichia coli*, *Lactococcus lactis*, *Pseudomonas aeruginosa*, *Sinorhizobium meliloti*, *Staphylococcus epidermidis*, *Streptococcus pneumoniae*, *Wolbachia pipientis*, and *Xanthomonas oryzae*. We performed joint clustering of intact genes and pseudogenes, to ensure that differences in how sequence clusters are defined did not influence the results. These 10 species varied widely in genome content and characteristics (**Extended Data Table 1**); for example, *Wolbachia pipientis* genomes encoded a mean of 897.0 intact genes (SD: 25.1) and 55.4 pseudogenes (SD: 20.8), while *Sinorhizobium meliloti* encoded a mean of 6032.8 intact genes (SD: 205.7) and 489.7 pseudogenes (SD: 53.4).

We separated gene/pseudogene clusters into three pangenome partitions, based on their frequency within a species: cloud (<=15%), shell (>15% and <95%), and soft-core (>=95%). We also further partitioned cloud clusters into ultra-rare, including clusters found in only one or two genomes (singletons and doubletons), and other-rare, referring to higher-frequency cloud clusters. Most pseudogene clusters were within the cloud partitions: mean of 95.46% (SD: 3.78%) vs. a mean of 84.01% (SD: 8.34%) for intact genes (**Extended Data Figure 1a**). This would be expected under the common assumption that pseudogenes are under relaxed purifying selection, such that they rapidly accumulate mutations which cause them to be split into separate sequence clusters. Some pseudogene clusters were in the soft-core partition (mean: 0.54%, SD: 0.66%), which often could not be annotated with Clusters of Orthologous Genes^11^ (COG) identifiers (**Extended Data Figure 1b**). For subsequent analyses we proceeded with COG-annotated clusters only (**Figure 1**).

**Figure 1:**
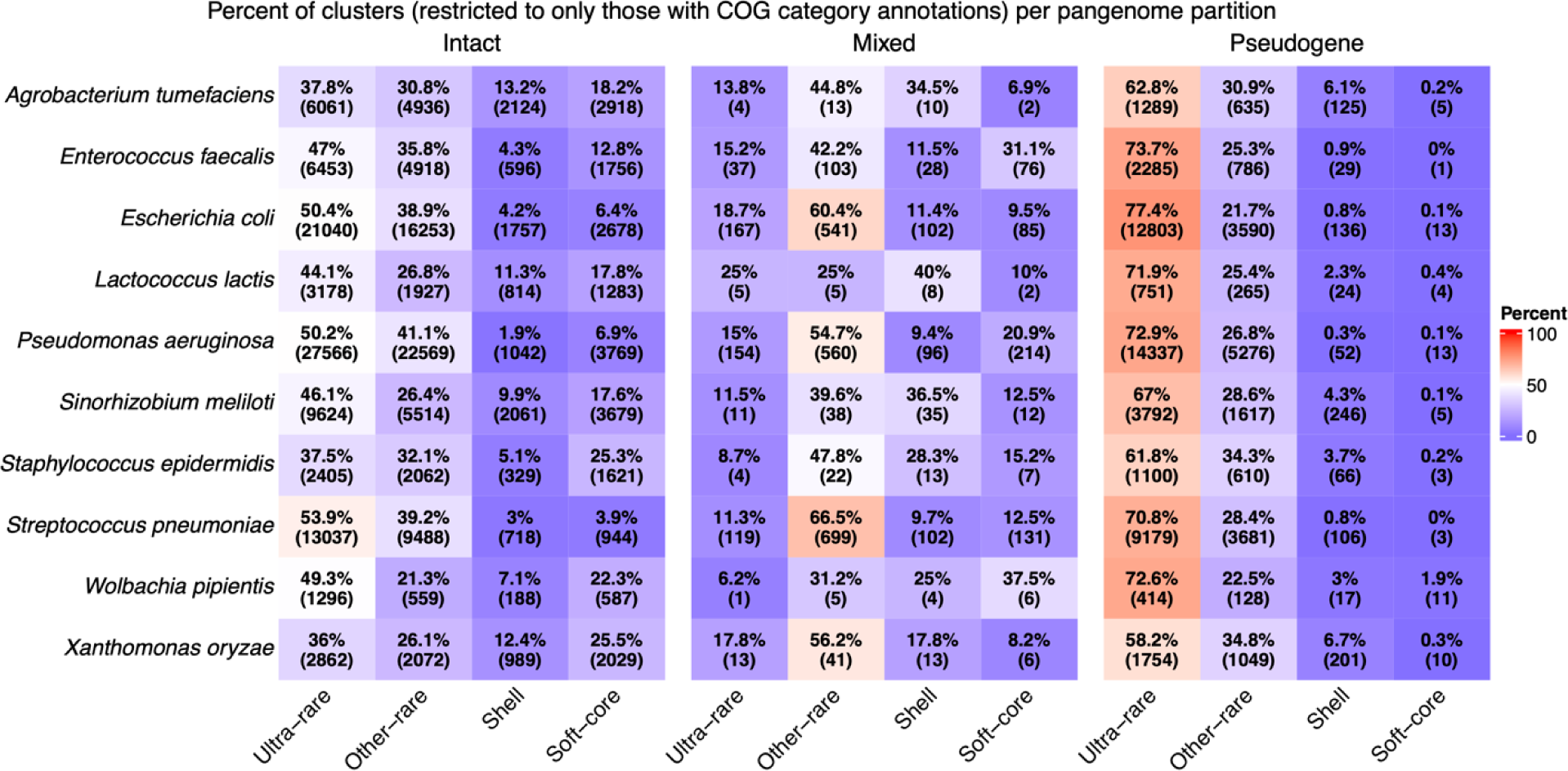
Distributions of gene or pseudogene sequence clusters by species and frequency in the pangenome, restricted to clusters that could be COG-annotated. Mixed elements are sequence clusters that include both pseudogenes and intact genes in the same cluster. Percentages correspond to the breakdown per species within a given element type (i.e. intact, mixed, or pseudogene) and raw counts are shown in parentheses.

We applied generalized linear mixed models, for each pangenome partition separately (excluding soft-core elements), to investigate which factors best explain whether a genetic element is an intact gene or a pseudogene. These models included 213,912, 3,650,010, and 12,234,597 elements for the ultra-rare, other-rare, and shell partitions, respectively. The fixed effects included each element’s COG functional category and whether the element was redundant with an intact gene with the same COG ID in the same genome. We included this ’redundancy’ effect because adaptive genes might neutrally degenerate only if they are complemented by an intact copy of the same gene family in the genome. The interaction between COG category and functional redundancy was also included as a fixed effect. Last, we also included species names, the interaction between COG category and species, and the interaction between functional redundancy and species random effects. All variables added significant information to the models, but there were some slight differences in their relative contributions. For instance, species identity and functional redundancy were particularly informative in the ultra-rare model compared to the more frequent categories of genes (**Extended Data Figure 2**), and certain species displayed different associations with pseudogenization by pangenome partition (**Extended Data Figure 3**).

We identified significant coefficients in the ultra-rare model (**Figure 2**), which provided insight into what factors were most associated with pseudogene status (*P* < 0.05). These coefficients represent decreased log-odds (logit) probabilities of an element being a pseudogene. Five COG categories were positively associated with pseudogenization: ‘energy production and conversion’ (C), ‘nucleotide transport and metabolism’ (F), ‘translation, ribosomal structure and biogenesis’ (J), ‘function unknown’ (S), and – most strongly – ‘mobilome: prophages, transposons’ (X). In contrast, ‘Cell cycle control, cell division, chromosome partitioning’ (D), was the sole COG category specifically associated with decreased pseudogenization. However, non-redundant elements were highly associated with decreased pseudogenization, over most COG categories. Non-redundant elements were also depleted for pseudogenes in the other-rare and shell models, but different COG categories were associated with pseudogenization overall (**Extended Data Figure 4**). The exception was an enrichment of pseudogenes in mobilome-associated elements in the other-rare partition.

**Figure 2:**
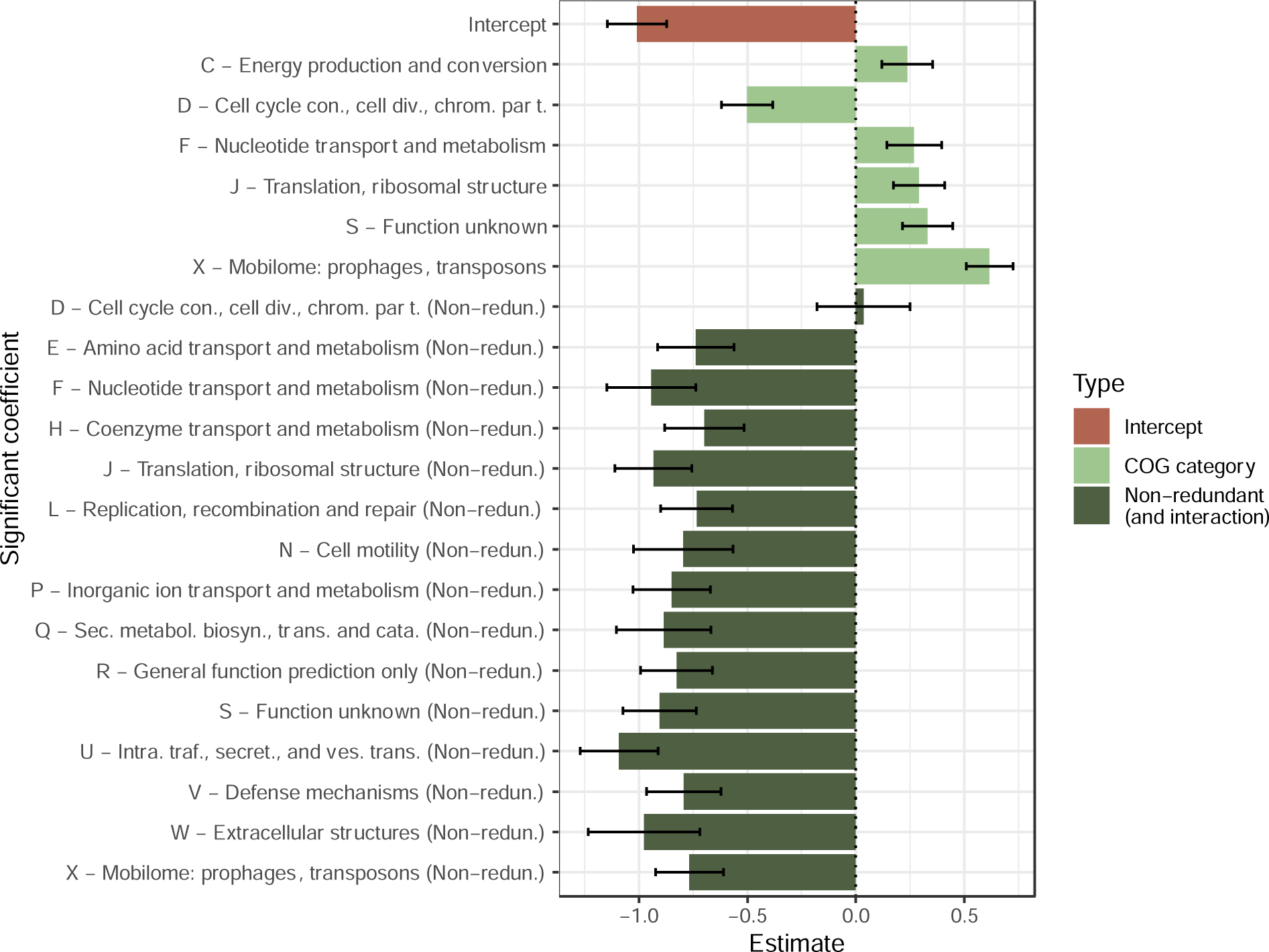
Summary of significant coefficients (*P* < 0.05) in generalized linear mixed model with singleton and doubleton (ultra-rare) element state (intact or pseudogene) as the response. This model was based on 213,912 separate elements. The predictors were each element’s annotated COG category, whether the element is redundant with an intact gene of the same COG ID (i.e. gene family, not COG category) in the same genome, and the interaction between these variables. The non-redundant coefficients represent the sum of the overall non-redundant coefficient and the interaction of non-redundancy and each COG category. Estimates correspond to logit (log-odds) values: estimates > 0 indicate an increased probability of an element being classified as a pseudogene. Error bars represent one standard error.

In the study of pangenome evolution, a key question is what proportion of rare genes are under selection or subject to genetic drift. This question is challenging to answer precisely; yet our models yield estimates of the percentage of genes found in functional categories depleted for pseudogenes, providing a lower bound for the percentage of adaptive genes. For instance, genes in COG category D and non-redundant genes in COG category E are two such pseudogene-depleted groupings. Based on these definitions, a mean of 19.41% (SD: 5.27%), 20.32% (SD: 6.84%), and 26.02% (SD: 7.05%) of intact genes are found in pseudogene-depleted groupings across the ultra-rare, other-rare, and shell partitions, respectively. The increasing percentage of genes classified as pseudogene-depleted as gene frequency increases from ultra-rare to shell is consistent with more frequent genes being more likely adaptive to their host. Nevertheless, an appreciable percentage (>19%) of ultra-rare genes are likely adaptive according to this estimate. Although non-redundancy was strongly and negatively associated with pseudogenization, only 24.39% of elements were non-redundant, which explains why only a minority of intact genes were categorized into pseudogene-depleted groupings. Conversely, 18.68% (SD: 5.62%), 13.29% (SD: 7.69%), and 3.65% (SD: 0.74%) of intact genes are found in groupings enriched for pseudogenes across these three partitions. The decreasing percentages as gene frequency increases is consistent with rarer genes being more likely deleterious to their host. Therefore, although rare accessory genes may on average be adaptive to their host genomes, a substantial fraction may also be deleterious. Most intact genes do not fall cleanly into either the pseudogene-enriched or -depleted category, meaning that these estimates represent rough lower bounds of how many genes are likely adaptive or deleterious.

Several COG categories were significantly enriched or depleted in pseudogenes, but these are broad functional groupings that can be difficult to biologically interpret. We investigated which individual COG IDs within significant COG categories were driving the overall signals in the ultra-rare model (see Online Methods). The clearest signal was of transposase-associated COGs being highly enriched among pseudogenes (mean of significant odds ratios: 5.10, SD: 6.86), which contrasted with other mobilome-associated COGs (**Extended Data Fig. 5**). We also identified several COGs highly associated with pseudogenization in specific species. For instance, anaerobic selenocysteine-containing dehydrogenases (COG0243, category C), were highly enriched for pseudogenes across multiple species, particularly in *Agrobacterium tumefaciens* (odds ratio: 103.6, *P* < 0.001). In addition, several COGs in category D involved in cell division and chromosome segregation were significantly depleted for pseudogenes, including BcsQ (COG1192), a ParA-like ATPase, which was significantly depleted for pseudogenes in six species (false discovery rate < 0.05).

This in-depth analysis of 10 species highlighted several functional categories enriched within rare pseudogenes, particularly for mobilome-related genes. Conversely, we identified a clear depletion of pseudogenes among non-redundant elements, which strongly suggests that even very rare accessory genes are often under selection to maintain a working copy in the genome. Taken together, these comparisons serve as a proof-of-concept that comparing extremely rare intact genes and pseudogenes can be useful for disentangling the action of evolutionary forces.

### Incorporating pseudogenes into comparisons of pangenome diversity and N_e_

We next investigated whether the inclusion of pseudogenes can help resolve the prior conflicting interpretations of the association between pangenome diversity and N_e_. To this end, we analysed 668 named prokaryotic species represented by at least nine genomes in the Genome Taxonomy Database^12^.

We first summarized pangenome diversity across these species. Species’ pangenome size and complexity have been characterised previously based on different metrics, including the mean number of genes per genome^2^ and genomic fluidity^6,13^. We computed these metrics for all species based on both intact genes and pseudogenes. In addition, as we were especially interested in rare elements, we computed the percentages of singleton genes and pseudogenes per species (i.e. those present in a single genome per species), based on repeated subsampling to nine genomes. Larger genomes tend to encode more singletons, both in mean number and percentage (**Extended Data Fig. 6a,b**). In addition, the percentage of intact singletons is highly correlated with genomic fluidity, but the traditional fluidity metric is sensitive to intermediate frequency accessory genes (**Extended Data Fig. 6c,d**), which can be driven by inconsistent species definitions or population structure within species. We therefore focused on the percentage of intact (s_i_) and pseudogene (s_p_) singletons for most analyses. All metrics ranged substantially across species for both intact genes (fluidity: 0.00-0.246; mean number: 836.4-8692.7; s_i_: 0.00-10.83%) and pseudogenes (fluidity: 0.014-0.513; mean number: 8.1-922.5; s_p_: 0.78-72.97%).

Values of s_i_ and s_p_ were positively correlated (Spearman’s ρ=0.57; *P* < 0.001), with deviations suggesting species-specific differences in selection on rare accessory genes (**Figure 3a**). For example, *Escherichia coli* has a relatively higher s_i_ value, consistent with selection to retain rare accessory genes, while the obligate intracellular bacteria *Chlamydia trachomatis* and *Rickettsia prowazekii* have lower values, suggesting less selective constraint on their rare genes. To summarize the s_i_ and s_p_ values per species, we focused on the s_i_/s_p_ ratio as a metric encompassing both intact gene and pseudogene pangenome diversity.

**Figure 3:**
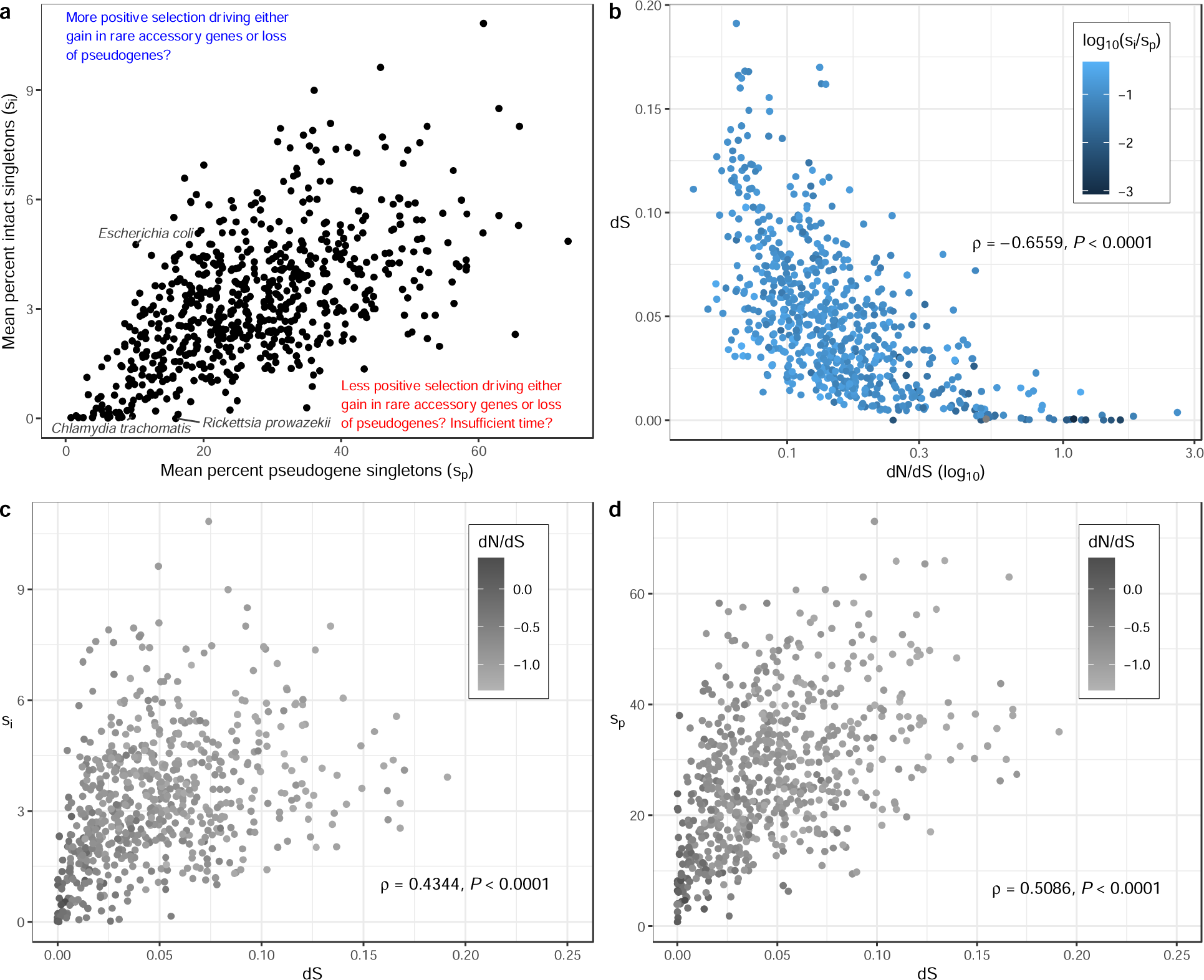
Distributions of singleton-based pangenome diversity and molecular evolution metrics. (a) Mean percentage of intact genes and pseudogenes that are singletons (i.e. genome-specific) per species. The mean percent singletons (for both intact genes and pseudogenes) per species was based on repeated subsampling to nine genomes (for up to 100 replicates). Possible (but non-exhaustive) drivers of higher or lower s_i_/s_p_ ratios are indicated alongside coloured arrows. Species mentioned in the main text are indicated. (b) Relationship between synonymous substitution rates (dS), a measure of strain divergence, and the ratio of the non-synonymous to synonymous substitution rates (dN/dS), coloured by s_i_/s_p_. Relationship between dS and (c) the mean percent intact singletons and (d) the mean percent pseudogene singletons, shaded by dN/dS. Across all panels, each point represents one of 668 prokaryotic species (>= 9 genomes each). Spearman correlation coefficients and *P*-values are indicated on panels b-d, and correspond to the comparison of the variables shown on the x and y axes.

For each species, we computed two proxies for N_e_: dN/dS and dS. These metrics were negatively correlated (**Figure 3b**), which could be due to the expected impact of N_e_ on each metric, and of course their direct dependence since dS is the denominator of dN/dS. However, the dependence structure may be more complex because recently diverged strains are biased towards higher dN/dS ratios due to insufficient time for purifying selection to purge deleterious non-synonymous mutations^14,15^. More generally, interpreting dS as N_e_ is questionable if there is widespread population substructure and uneven sampling of sequenced strains (although this has been debated: see Discussion). For this reason, we focused on dN/dS as an inversely related proxy for N_e_, and we considered dS as a measure of the divergence between the subset of analysed genomes per species, but not necessarily as representative of species-wide N_e_. Under this interpretation, the observed positive correlation between dS and both s_i_ and s_p_ (**Figure 3c and 3d**) is expected simply because these are all measures of genome divergence.

We next recapitulated the previously observed association between dN/dS and standard measures of pangenome diversity^2,3^, and then explored whether dN/dS is also associated with s_i_/s_p_. We found that the mean number of genes per species was not significantly associated with dN/dS (**Figure 4a**), but both genomic fluidity, and s_i_ were significantly negatively correlated (**Figure 4b and c**; Spearman correlations, *P* < 0.05). Although the mean number of genes has been considered in this context previously, it is a measure of overall pangenome size, and is not a direct measure of gene content diversity, which could explain why we did not observe a significant relationship. In contrast, the latter two observations agree with past work^2,3^, but, as discussed above, the biological interpretation of these associations is unclear. Notably, we also found s_i_/s_p_ to be negatively associated with dN/dS (Spearman’s ρ=-0.22; *P* < 0.001; **Figure 4d**), although less strongly than s_i_ alone (Spearman’s ρ=-0.54; *P* < 0.001). These results were qualitatively robust to the number of genomes subset when computing s_i_ and s_p_ (**Extended Data Figure 7**). In addition, although dS was significantly associated with s_i_ (**Figure 3c**), it was not significantly associated with s_i_/s_p_ (Spearman ρ=0.07; *P*=0.08). Taken together, these results highlight that s_i_ remains associated with dN/dS even after normalization by s_p,_ but that the association with dS is lost after this normalization. If pseudogene presence/absence diversity is assumed to be a proxy for neutral gene content diversity, this finding suggests that intact singleton gene prevalence is particularly associated with selection efficacy (dN/dS), and not simply with strain divergence. These results are consistent with s_i_/s_p_ behaving somewhat analogously to dN/dS as a measure of the efficacy of selection. As a higher fraction of rare genes (relative to pseudogenes) are retained when selection is more effective, this is consistent with many singleton genes conferring adaptive benefits, and/or some singleton pseudogenes being slightly deleterious.

**Figure 4:**
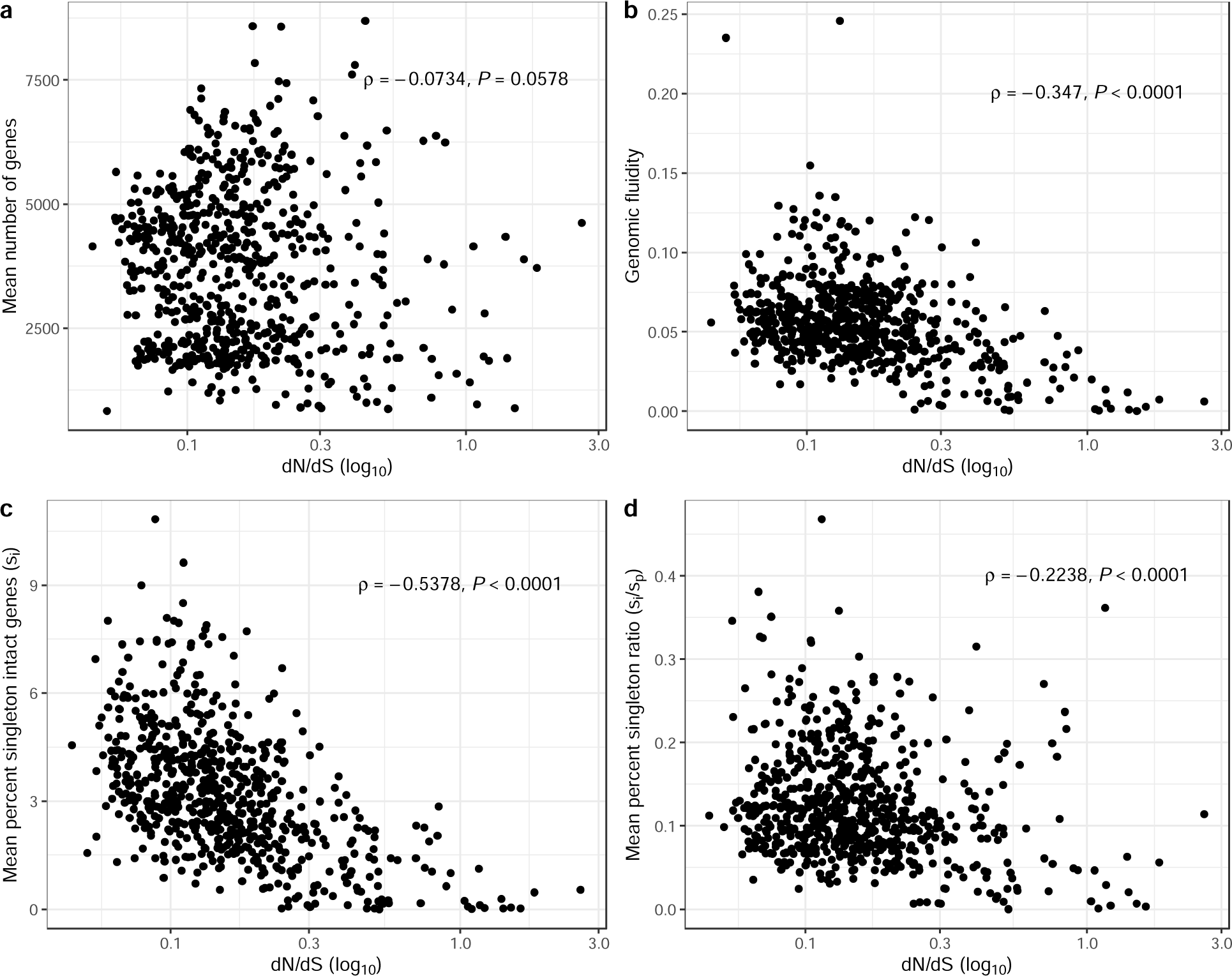
Associations between pangenome diversity metrics and estimated efficacy of selection (dN/dS). Each panel presents the association between the ratio of non-synonymous to synonymous substitution rates (dN/dS; across each species’ core genome), plotted on a log_10_ scale, and one of the following measures: (a) the mean number of genes per genome, (b) genomic fluidity, (c) the mean percent of intact singletons, and the percentage of singleton intact genes normalized by the percentage of singleton pseudogenes per species. Each point is one of 668 prokaryotic species. The Spearman correlation coefficients and *P*-values are indicated.

There are certain species included in our analyses that are outliers with low values of dS and/or high values of dN/dS (**Figure 3b**), which could potentially impact these correlations. In addition, within-species dN/dS systematically varies with dS, which could also impact our conclusions. To address these issues, we re-ran our analysis with outliers removed and using partial Spearman correlations that control for dS (**Figure 5**). With minor exceptions, the results remained qualitatively unchanged, which demonstrates that these factors are not driving the signal.

**Figure 5:**
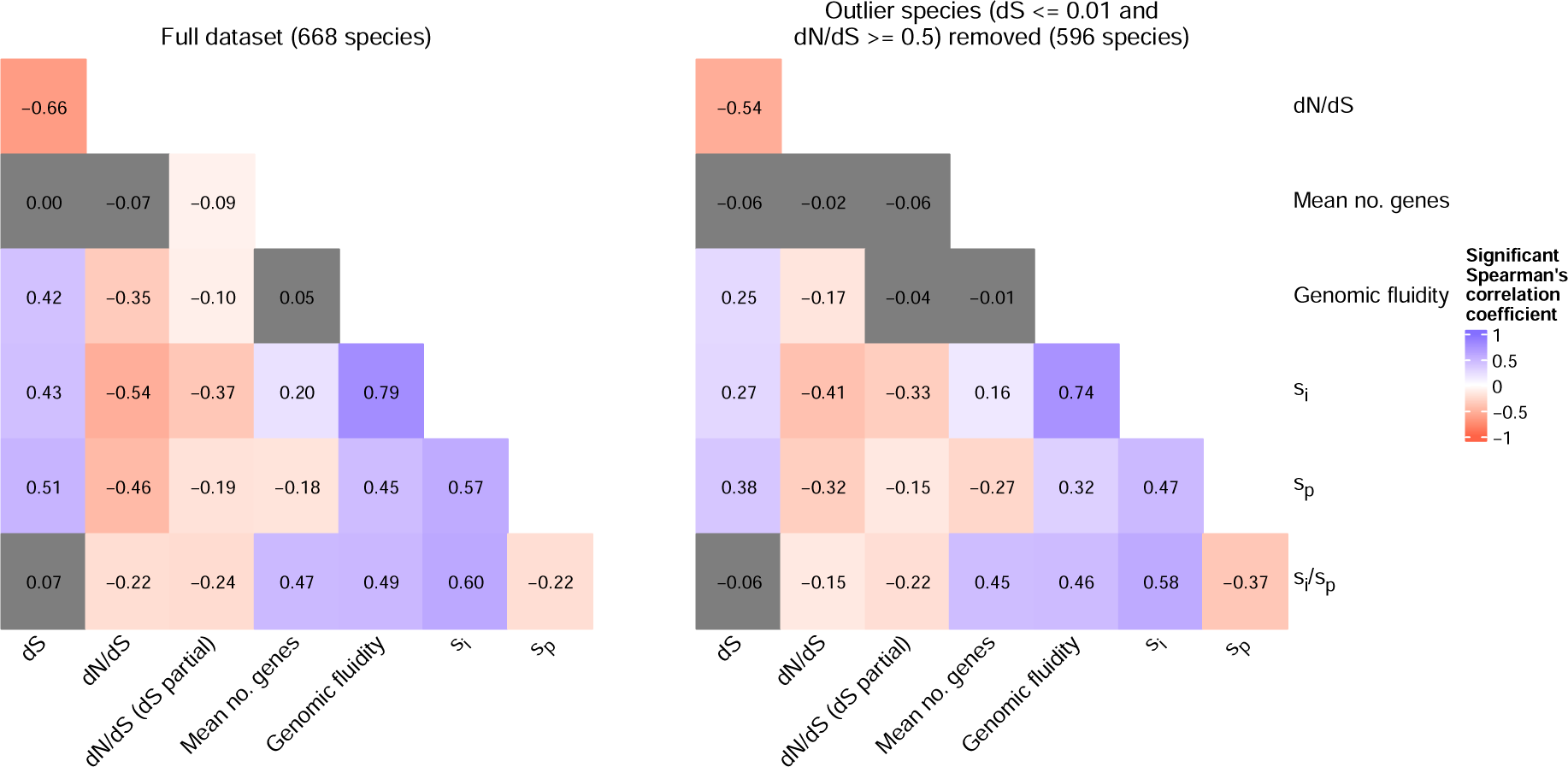
Spearman’s correlations between molecular evolution and pangenome diversity metrics. Each cell represents the correlation coefficient for a pairwise comparison of variables. Coloured cells are significant (*P* < 0.05), and non-significant cells are dark grey. The left plot includes all species while the right plot provides the results based on a subset of species, with outliers for dS and dN/dS removed. The column dN/dS (dS partial) corresponds to partial Spearman correlations between dN/dS and each variable, controlling for dS.

Another potential issue is that the species included in our analyses, although they span substantial prokaryotic diversity, were biased towards specific groups, particularly Gammaproteobacteria (286 species) and Bacilli (161 species). As there is substantial variation in pangenome diversity and evolutionary metrics at the class level for the species we considered (**Extended Data Figure 8**), taxonomic biases could potentially be driving the correlations with s_i_/s_p_ we observed. To account for this, we conducted a linear modelling analysis, where a separate model was generated with each of the four pangenome diversity measures as the response, and dS, dN/dS, and taxonomic class as predictors. All models were highly significant (*P*<0.001; **Extended Data Figure 9**) and ranged in adjusted R^2^ values from 0.197 to 0.420 for the s_i_/s_p_ and s_i_ models, respectively. All but one class (Bacilli) were significant predictors in at least one model, and the classes Clostridia, Bacteroidia, and Chlamydiia were significant predictors across all four models. Similarly, dS was a significant predictor of all pangenome diversity metrics except for s_i_/s_p_. In contrast, dN/dS was a significant predictor for all pangenome diversity metrics except for the mean number of genes. This analysis demonstrated that, despite class-specific differences in pangenome diversity, our overall inferences are robust to taxonomic class as a confounder.

A final caveat of these analyses is that higher s_i_/s_p_ values could be explained by selection to preserve rare intact genes or to purge rare pseudogenes. If there were selection for pseudogene loss, then the pseudogene content per genome would be expected to be lower in species with higher selection efficacy. Contrary to this prediction, the mean percent of species’ genomes covered by pseudogenes was not significantly associated with dN/dS (Spearman’s ρ = 0.063; *P* = 0.104; **Figure 6a**), which is inconsistent with a model of widespread slightly deleterious pseudogenes that are purged only in species with sufficiently high N_e_. However, pseudogene coverage is negatively but weakly associated with dS (Spearman’s ρ = -0.090; *P* = 0.020; **Figure 6b**), which highlights that this interpretation is dependent on the assumption that dN/dS is a more appropriate proxy than dS for N_e_ across these genomes (see Discussion). Together, the lack of association between pseudogene content and dN/dS, and the weak association with dS, argue against adaptive purging of rare pseudogenes as a major driver of variation in s_i_/s_p_.

**Figure 6:**
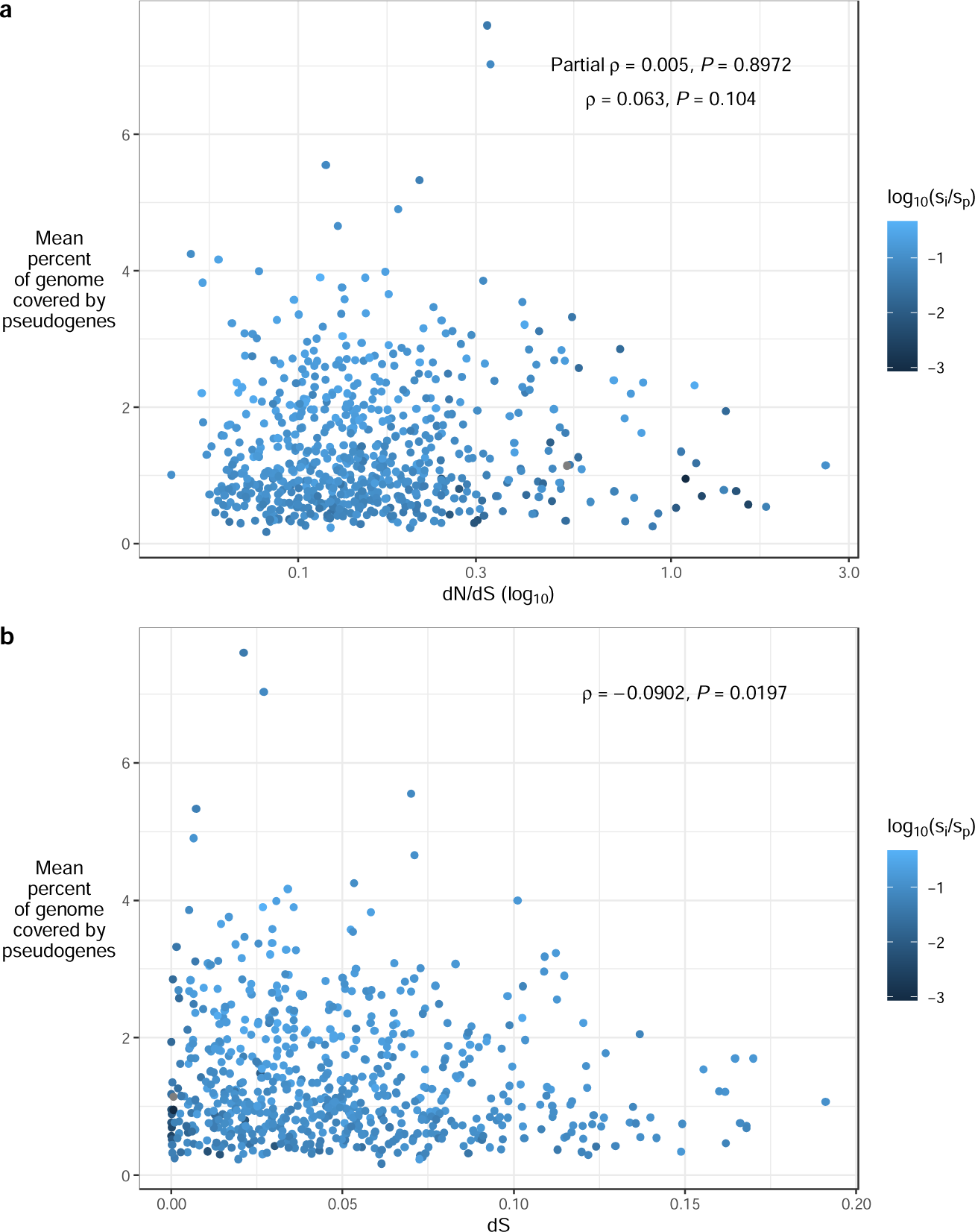
Weak relationships between the 312 mean percent of each species’ genome covered by pseudogenes and the (a) within-species ratio of the non-synonymous to synonymous substitution rates (dN/dS) and (b) the within-species synonymous substitution rate (dS). Each point corresponds to one of 668 species. Values of si/sp are overlaid on a log10 scale on both panels. The result of Spearman correlation tests between the variables plotted on the x and y are indicated (including the partial Spearman correlation controlling for dS in panel a).

## Discussion

The ability to distinguish neutral and adaptive models of pangenome evolution has been hindered by a lack of tools to test for selection acting on gene content. This contrasts with an established toolkit of tests for selection at the nucleotide and protein levels, including dN/dS and its extensions. Here we propose pseudogenes as a reference for distinguishing neutral and adaptive forces acting on pangenomes – particularly rare genes. We show that the association between pangenome diversity and synonymous-site variation disappears after correcting for pseudogene diversity with the s_i_/s_p_ metric, while the association with dN/dS is maintained. This indicates that a higher proportion of intact singleton genes (relative to singleton pseudogenes) are present when selection is more effective. This is consistent with many rare intact genes, but not all, conferring host-adaptive functions. These genes are more likely to be retained when selection is efficient^3^ (such as in *E. coli*), and more likely to degenerate neutrally and become pseudogenes in species with lower N_e_ (such as obligate intracellular bacteria). Our results could also be explained by widespread slightly deleterious rare pseudogenes, which can be purged only in species with high N_e_, but we did not detect a significant association between dN/dS and pseudogene content (and only a weak association with dS), making this less likely.

A common explanation for widespread selection on rare accessory genes is adaptation to highly specialized niches^16–18^. While genes recently acquired through horizontal gene transfer are often hypothesised to confer niche-specific adaptations^4^, it is challenging to make high-confidence inferences without knowing the background of all recently transferred genes that were not retained – and are thus, by definition, unobservable. By focusing on pseudogenes, which are observable but likely to evolve mostly by drift, we can establish a (nearly) neutral background against which to discern potentially niche-specific adaptations.

We relied on the assumption that any selection pressures acting upon pseudogenes tend to be of much lower magnitude compared to intact genes. In other words, we assumed that, overall, the pseudogene instances we identified do not reflect adaptive gene loss^19^ (which is unlikely to substantially increase with selection efficacy, as described above), nor do they contain beneficial regulatory sequences for modulating gene expression levels^20^. This second possibility would be inconsistent with the positive association we observed between s_i_/s_p_ and selection efficacy. Instead, our results are consistent with rare pseudogenes evolving under a regime closer to neutrality relative to rare intact genes.

Our enrichment test results highlight that a significant proportion of rare accessory genes are under selection to be retained. Notably, 19% of ultra-rare intact genes are in COG categories significantly depleted for pseudogenes. We stress that this is a rough approximation and does not imply that precisely 19% of ultra-rare intact genes have adaptive value. We hypothesise that many such genes are under effective purifying selection, while relaxed purifying selection could account for the observed enrichment of transposons among pseudogenes. Similarly, the enrichment of selenocysteine-containing dehydrogenases among pseudogenes could similarly reflect relaxed or sporadic purifying selection on these elements, which is interesting as selenium, selenocysteine’s defining component, is sporadically used across the prokaryotic tree^21^.

Gene-level selection could also account for certain observations. For instance, the DNA partitioning protein highly enriched in intact ultra-rare genes, COG1192, is a known plasmid-encoded element predicted to be involved in plasmid partitioning^22^. There could be an ascertainment bias toward identifying such genes as intact, because were they pseudogenized or lost the entire plasmid might not be transferred to daughter cells. Similar biases could also account for why prophage and plasmid-associated elements in the mobilome more generally are depleted among pseudogenes, although these elements are also more likely to be adaptive to the host genome^23,24^.

Pseudogene diversity can be influenced by many factors, including life history. For instance, obligate intracellular bacteria are characterized by widespread degeneration of their genome, followed by streamlining^25^. Depending on a species’ stage in this evolutionary process, its genome could be enriched or depleted for pseudogenes relative to other bacteria. This likely accounts for certain s_i_/s_p_ outliers, such as the obligate intracellular bacteria *Rickettsia prowazekii*, which had the lowest s_i_/s_p_ ratio. Accordingly, our framework could be improved by incorporating per-species parameters of pseudogene gain and loss dynamics.

Last, an important assumption underlying these results is that dN/dS is an accurate proxy for selection efficacy, and thus N_e_, while dS does not represent species N_e_. This is counterintuitive, as nucleotide diversity at neutral sites is the standard metric for calculating N_e_. However, the validity of this metric in prokaryotes has been debated^26,27^. The estimated dS values would only be generalizable to the overall species (i.e., including all unsampled genomes) if this sampling is representative of the actual diversity in nature. For instance, if sequenced genomes are more likely to represent strains adapted to distinct local environments, and be depleted for near-identical strains from the same environment, then this would not be representative. It is highly unlikely, particularly given the low numbers of genomes considered, for the strains considered in these analyses to accurately reflect the strain diversity and population substructure across species. Instead, we believe dS is more appropriately considered a measure of divergence time among the subset of genomes analysed, but not generalizable across the entire species. In contrast, dN/dS across core genes is more appropriate to generalize across a species when calculated based on a subset of genomes, as this is expected to be similar for all pairwise strain comparisons (albeit with variation depending on strain divergence time), regardless of whether the genomes are representative of the overall species’ strain diversity. However, dN/dS is also an imperfect measure, particularly because synonymous sites do not evolve completely neutrally^28^. Regardless, it is uncontroversial to assume that selection acting on synonymous sites is, on average, much weaker compared to on non-synonymous sites. Accordingly, although dN/dS may be inappropriate to use for explicitly calculating N_e_, the species’ relative ranks for this measure will correspond inversely to their relative N_e_ rank, all else being equal.

Despite these caveats, our work highlights the value of using pseudogene diversity as a neutral null^29^ for evaluating the evolutionary forces acting upon intact accessory genes. Establishing true neutrality in microbial genomes is challenging^30^, but the clear association we identified between dN/dS and s_i_/s_p_ suggests that pseudogene presence/absence diversity can provide insight into how rare accessory genes evolve. Crucially, rare genes in nearly all functional categories are less likely to be pseudogenes when there is no redundant gene copy in the same genome, indicating that even very rare accessory genes are commonly under selection to maintain an intact copy in the genome. Using this pseudogene-based comparative approach, we show that a neutral pangenome model can be rejected and identify which types of rare genes, based on their functional annotation and which species encode them, are more likely to be retained.

## Code and data availability

The code used for the analyses in this manuscript is located at https://github.com/gavinmdouglas/pangenome_pseudogene_null and the key data files are available on Zenodo (DOI: 10.5281/zenodo.8326664). All analysed genomes are publicly available as part of NCBI RefSeq/GenBank.

## Acknowledgements

We would like to thank Ford Doolittle for providing motivating ideas, and for advice and feedback throughout this project. We would also like to thank Louis-Marie Bobay for reading a draft of this manuscript and providing feedback, and Adam Eyre-Walker for providing constructive comments. GMD is supported by a Natural Sciences and Engineering Research Council of Canada (NSERC) Postdoctoral Fellowship and BJS is supported by an NSERC Discovery Grant.

## Ethics declarations

The authors declare that they have no competing interests related to the content of this article.

## Online Methods

### Dataset processing – In-depth pangenome analysis

We conducted an analysis of 10 bacterial species with a relatively high number of genomes (ranging from 135-6,916). We selected these species from the set identified for the broad pangenome analysis (see below), but that were also represented by > 100 genomes that were not phylogenetically redundant. For these data, we clustered both intact genes and pseudogenes with cd-hit^31^ version 4.8.1 with an identity cut-off of 95% over at least 90% of both compared sequences. This clustering was performed on all genes and pseudogenes across all ten species. We assigned clusters to pangenome partitions as described in the main text. Note that we defined the ultra-rare partition to include doubletons, and not only singletons, to account for cases where two highly similar strains are present with the same ultra-rare gene. As there were > 100 genomes considered for each species within this analysis, doubletons also correspond to highly rare genes.

We functionally annotated each resulting cluster with COG IDs and categories^11^ using eggNOG-mapper^32^ version 2.1.6 (based on eggNOG orthology data^33^ version 5.0.2) with DIAMOND^34^ version 2.0.14 and these parameter options: --score 60, --pident 40, --query_cover 20, --subject_cover 20, --tax_scope auto, and --target_orthologs all. This was performed for individual elements separately (i.e. the original sequences rather than the cluster representatives), and for database sequence matches to pseudogene hits. We focused on the database sequence matches for pseudogene hits, as eggNOG-mapper annotates protein sequences, which is problematic for most pseudogenes as the protein-coding information is generally lost.

Accordingly, annotating the corresponding database hits per pseudogene is a more reliable way of assigning putative function. We used majority rule of all member sequences per cluster to assign individual COG IDs and categories, and the same approach for assigning functions to individual pseudogene sequences based on database sequence annotations. We assigned COG categories based on a mapping of COG IDs from the COG 2020 database release. This was performed as the raw output COG categories were based on an earlier version of the database that did not include mobilome (category X) annotations.

### Generalized linear mixed models

Generalized linear mixed models were fit in R using the glmmTMB^35^ package v1.1.5, one for the ultra-rare, other-rare, and shell pangenome partitions, respectively. Only COG-annotated elements were included in these models, excluding those annotated by the (rare) A, B, Y, and Z COG categories only. We used the binomial family and nlminb optimization algorithm with 1000 set for both iter.max and eval.max. The full R-style formula for each model was:

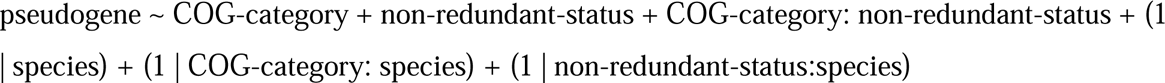

pseudogene ∼ COG-category + non-redundant-status + COG-category: non-redundant-status + (1 | species) + (1 | COG-category: species) + (1 | non-redundant-status:species)

In this formula, random effects are specified as those in parentheses including “1|” and interaction terms are indicated with “:”. The response was a Boolean variable indicating whether each element is a pseudogene. The COG-category variable is categorical indicating the one-letter COG category code that each element belongs to. In cases where elements were members of multiple categories, duplicate rows were created for each category. The Transcription category (K) was selected as the first level, to be used for the intercept, as it was the most consistently abundant COG category across all three partitions (third in the other-rare and shell, and fourth in ultra-rare). The non-redundant-status variable was a Boolean variable indicating whether each element was not redundant with another intact element of the same COG ID (gene family, not category) in the same genome. This negative formulation of redundancy (i.e. whether an element is not redundant, rather than whether it is redundant) was chosen as most elements were redundant, and so we decided to set the default level in each model (False) to be more representative. The species variable corresponded to the name of the species encoding each element.

We also fit simpler models with subsets of these variables and computed Akaike Information Criterion (AIC) values for each model, that allowed us to compare across models and investigate whether more complex models provide significantly more information. We visualized the AICs per model based on normalized scores that transformed the minimum model AIC per partition to be 0 and the maximum model AIC per partition to be 1.

Finally, for each significant COG category in the ultra-rare generalized linear model (excluding those interacting with non-redundancy), we systematically tested whether individual COG IDs were enriched for pseudogenes based on Fisher’s exact tests comparing the number of pseudogene and intact genes within each COG ID (and with the same redundancy status and in the same species) compared to the background of all other elements with the same redundancy status in the same species.

### Dataset processing – broad pangenome analysis

We downloaded all genomes used in this study from the Genome Taxonomy Database^12^ release 202. We identified all species in this database with at least ten high quality genomes, based on these criteria: (1) marked as passing the minimum information about a metagenome-assembled genome^36^ check; (2) CheckM^37^ completeness > 98% and contamination < 1%; (3) fewer than 1000 contigs; (4) contig N50 > 5000; (6) fewer than 100,000 ambiguous bases. We also restricted our analyses to genomes in RefSeq (rather than those in GenBank only), except for *Wolbachia pipientis* genomes, which were numerous but primarily limited to GenBank. For species with more than twenty genomes, we randomly sampled down to twenty genomes. We identified 670 species that fit these criteria and downloaded the corresponding genomes. Certain genomes had been relabelled or removed from NCBI since the release of Genome Taxonomy Database release 202, which resulted in a minimum of nine genomes per species (we eliminated two species with fewer than nine genomes). We annotated all genomes with Prokka^38^ version 1.14.5 with the –kingdom, --compliant, and –rfam options. We also specified the —metagenome flag for all genomes with 50 or more contigs. We ran Panaroo^39^ version 1.3.0 on all output GFFs, with the –remove-invalid-genes and --clean-mode strict options. We then ran Pseudofinder^40^ on the Prokka-output GenBank files to identify all putative pseudogenes, using protein sequences from the UniRef90 database^41^ (UniProt KB release 2022_01) as a reference database. We restricted the output to intergenic pseudogenes specifically, as the other pseudogene types identified by Pseudofinder correspond to divergent intact coding sequences (in length or modularity), which are difficult to interpret as truly degenerating sequences, and could simply represent functionally divergent proteins. We performed three filtering steps on the output intergenic pseudogenes. Specifically, we excluded all (1) pseudogene calls within 500 bp of contig ends, (2) pseudogenes of called length < 100 bp or > 5000 bp, and (3) pseudogenes that substantially differed from the mean size of all matching database hits (mean database size – pseudogene size was inclusively required to be between -500 bp and 2000 bp). Pseudogenes were clustered with cd-hit using the same settings as described above. Where possible, these commands were parallelized with GNU Parallel^42^ version 20161222.

### Pangenome metric computation

The mean numbers of singletons (whether of intact genes or pseudogenes) per species were identified after repeated subsampling to nine strains per species and then comparing the overlapping genes/pseudogenes. This procedure was repeated for up to 100 replicates (or until the maximum number of strain combinations was reached) and the number of singletons per genome was computed across all replicates. Note that for a supplementary analysis this subsampling was also conducted for subsamples of three and 20 genomes.

the estimated number of singleton genes for a given species, given a number of subsampled genomes *k*, *U_k_* is defined as: 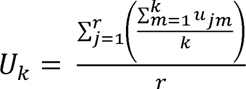, where *r* is the total number of subsampled replicates and *u_jm_* is the number of genome-specific genes found in genome *m* (based on genomes subsampled in replicate *j*). If there are *N* genomes in total for a given species, then 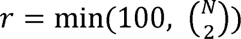. The mean percentage of intact gene singletons per species can then be calculated as: 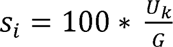, where *G* is the mean number of genes per genome (across all *N* genomes). This same procedure was repeated for pseudogenes, except that the numbers of singleton pseudogenes were computed per subsample replicate, and the mean percentage of pseudogenes per species (s_p_) was calculated based on the average number of pseudogenes per genome. To be clear, this formulation means that the s_i_/s_p_ metric corresponds to a comparison of the percentage of singleton intact and pseudogene calls overall per species, rather than of calls within each individual genome.

### Evolutionary metric computation

We performed codon-aware multiple-sequence alignment of all ubiquitous and single-copy genes sequences per-species with muscle^43^ version 3.8.1551, based on the HyPhy^44^ version 2.5.36 codon-aware workflow (https://github.com/veg/hyphy-analyses/tree/master/codon-msa). We then concatenated the core gene alignments per species with a Python script (cat_core_genome_msa.py) and computed pairwise dN/dS and dS for each combination of strain pairs per species with an additional script (mean_pairwise_dnds.py). Both scripts, and the bash commands for running the codon-aware alignments, are available in v1.1.0 of this repository: https://github.com/gavinmdouglas/handy_pop_gen. The latter script identifies potential non-synonymous and synonymous mutation sites between each sequence pair using the NG86 approach^45^. We computed the mean values across all pairwise strain comparisons, resulting in a single measure of dN/dS and dS per species.

### Linear models

We built linear models using the lm function in R to predict pangenome diversity, based on (per species) either the mean number of genes, the genomic fluidity, s_i_, or s_i_/s_p_. The predictors included dS, dN/dS, and taxonomic class. Classes with <= 5 member species were collapsed into the “Other” category, which was set as the intercept for the models. One species, *Rickettsia prowazekii*, was excluded from this analysis due to values of zero for s_i_ and s_i_/s_p_. We transformed all continuous variables to be normally distributed, except for the mean number of genes, which was already normally distributed. We performed a square-root transformation of the genomic fluidity, s_i_, s_i_/s_p_, and dS values. The dN/dS values were especially right skewed and required a negative inverse transformation (-1 * 1/x, where x is each dN/dS value) to be normalized. We then converted each continuous variable to standardized units, by mean-centring and dividing by the standard deviation. This step means that the model outputs refer to units of standard deviation per variable, which makes it possible to compare the magnitude of coefficients across models with different response variables.

### General analyses

No tests for statistical power were conducted to determine the sample sizes required for this study, but we used genomes from all available species in the Genome Taxonomy Database of sufficient quality. All analyses were conducted in R v4.2.2. Figures were generated with ggplot2^46^ v3.4.0, with the exception of the heatmaps, which were created with the ComplexHeatmap^47^ package v2.14.0.

