## Supplementary Information for "Pseudogenes as a neutral reference for detecting selection in prokaryotic pangenomes"

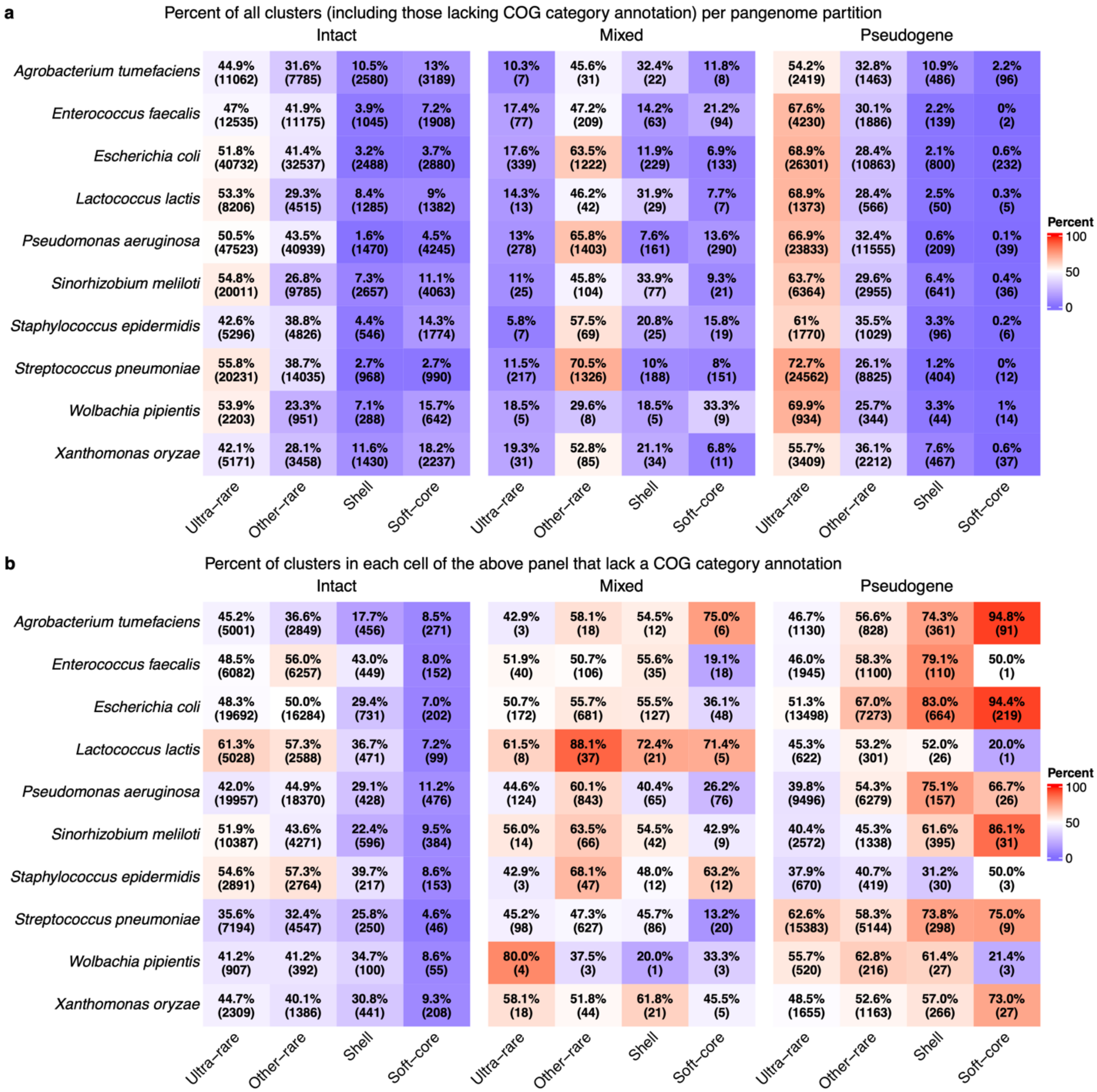


**Extended Data Figure 1**: Frequency distributions of clusters by species, pangenome partitions, and element type. Mixed elements are those that include both pseudogene and intact gene sequences in the same cluster. (a) Distribution for all clusters, including those that could not be annotated with a Clusters of Orthologous Genes (COG) identifier. Percentages correspond to the breakdown per species within a given element type (i.e. intact, mixed, or pseudogene). (b) Breakdown of the numbers and percentages of clusters that could not be COG annotated (i.e. the percentages correspond to the how many of the clusters per cell in panel a could not be COG annotated).

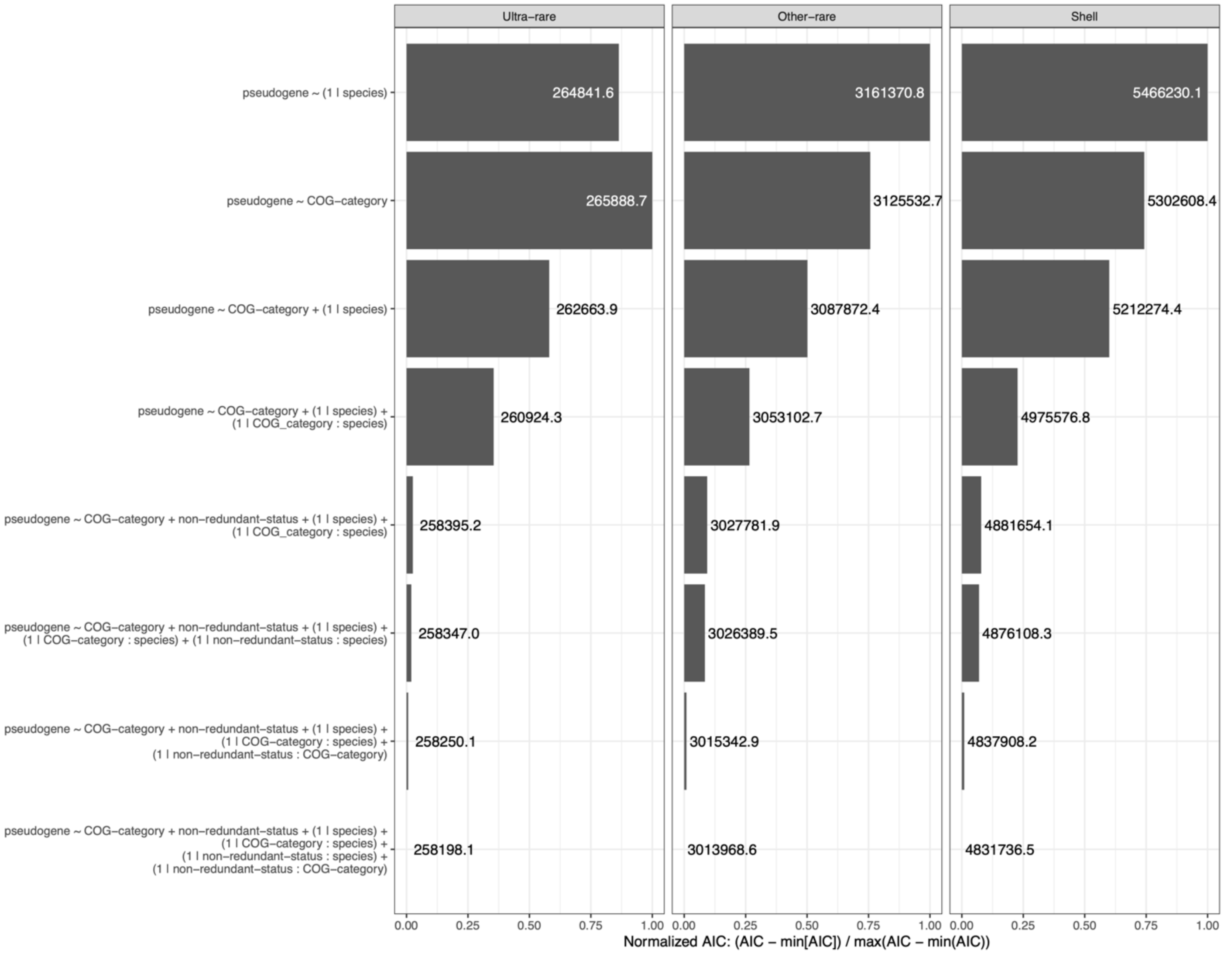


**Extended Data Figure 2**: Akaike Information Criterion (AIC) values for each generalized linear mixed model across the three tested pangenome partitions. These partitions contained 213,912, 3,650,010, and 12,234,597 separate elements for the ultra-rare, other-rare, and shell partitions, respectively. The model formulas are indicated on the y-axis (see Online Methods for explanation). AIC values were normalized to range from 0-1 for better visualization of relative differences, and the raw AIC is indicated beside each bar.

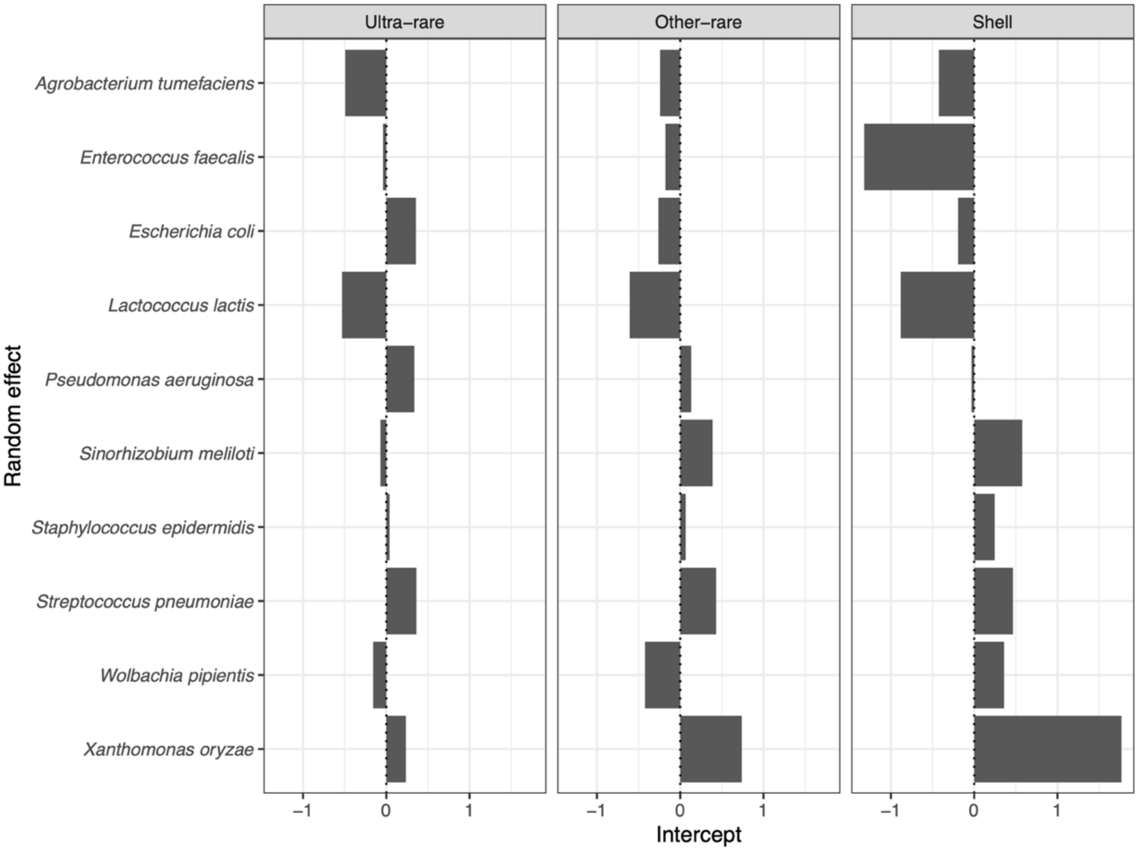


**Extended Data Figure 3**: Intercepts for each species random effect level in the generalized linear mixed models. Each model is labelled by the corresponding pangenome partition. These partitions contained 213,912, 3,650,010, and 12,234,597 separate elements for the ultra-rare, other-rare, and shell partitions, respectively. Estimates correspond to logit (log-odds) values: estimates > 0 indicate an increased probability of an element being classified as a pseudogene. Note that each model has a different overall intercept value, meaning that the relative differences in species per model is most informative to compare across models, rather than the absolute intercept values (for instance, *Enterococcus faecalis* has the highest magnitude negative intercept in the shell model, but is of relatively low magnitude in the other models).

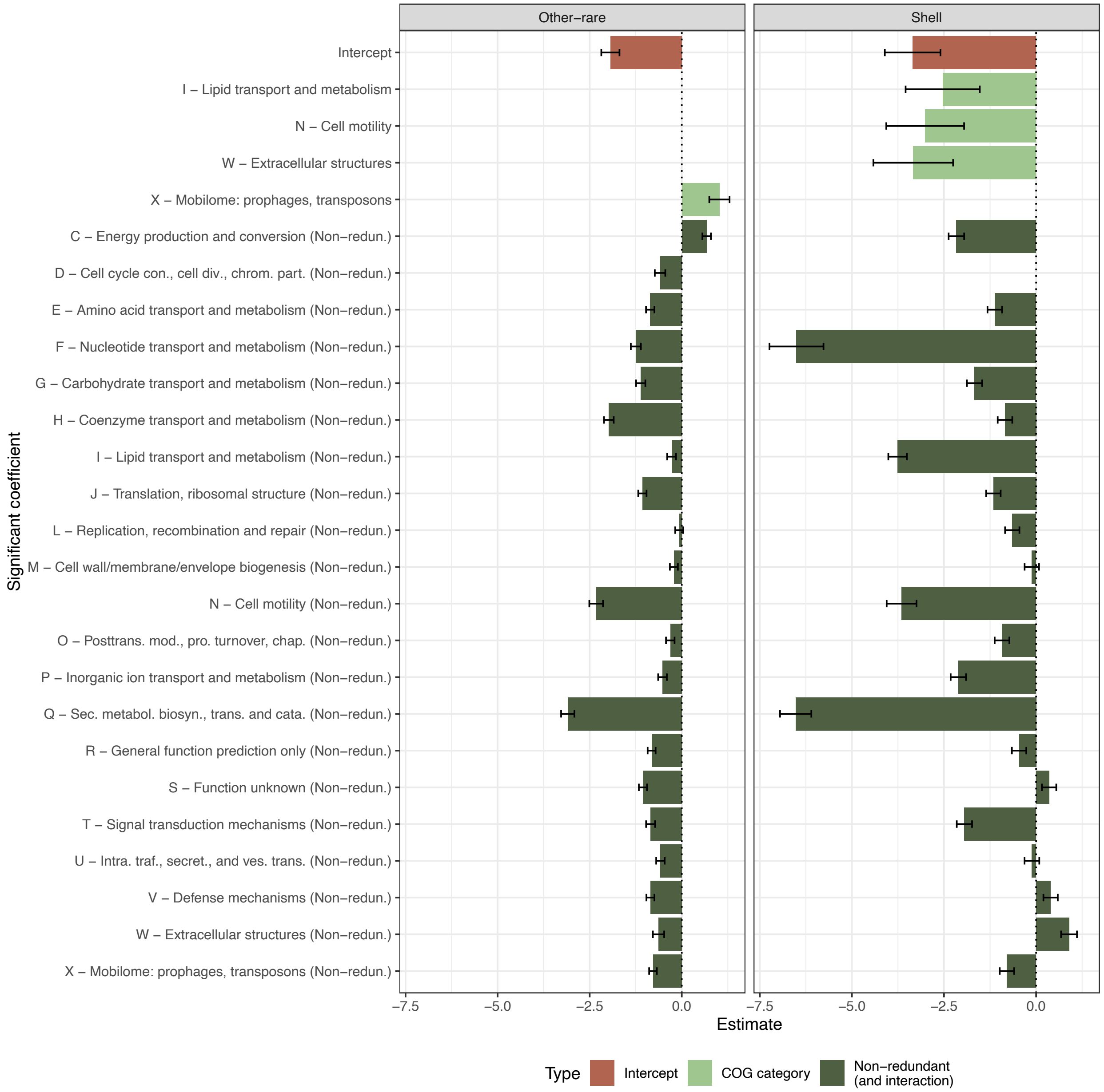


**Extended Data Figure 4**: Summary of significant coefficients ($P < 0.05$) in generalized linear mixed models fit to other-rare and shell pangenome partition elements, which corresponded to 3,650,010 and 12,234,597 separate elements, respectively. Model response was element state (intact or pseudogene). The predictors were each element's annotated Clusters of Orthologous Genes (COG) category, whether the element is redundant with an intact gene of the same COG ID (i.e. gene family, not COG category) in the same genome, and the interaction between these variables. The non-redundant coefficients represent the sum of the overall non-redundant coefficient and the interaction of non-redundancy and each COG category. Estimates correspond to logit (log-odds) values: estimates > 0 indicate an increased probability of an element being classified as a pseudogene. Error bars represent one standard error.

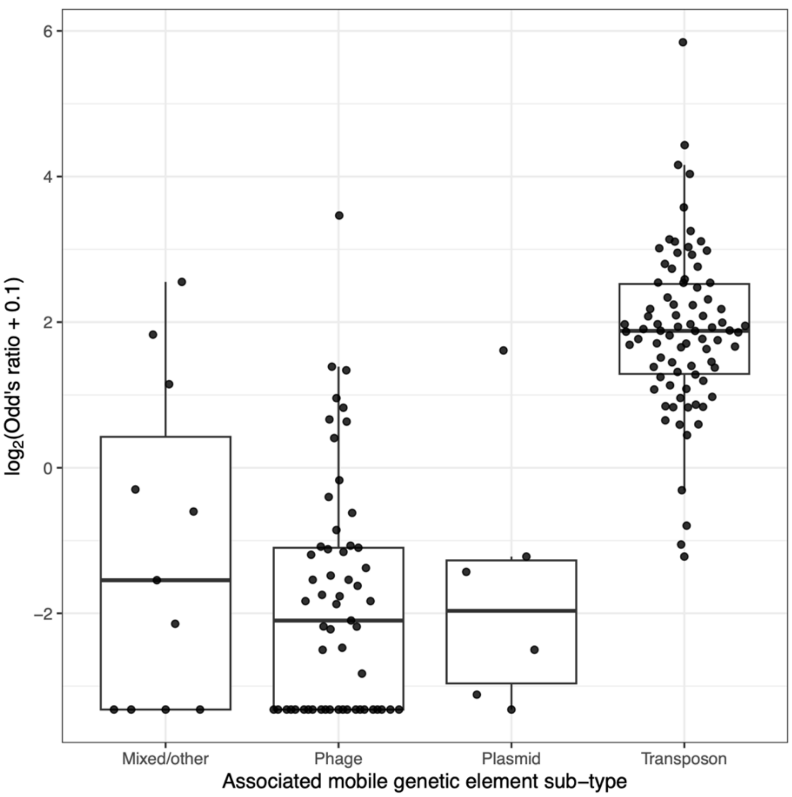


**Extended Data Figure 5**: Log-odds ratios for the 156 significant (Fisher's exact test, false discovery rate < 0.05) Clusters of Orthologous Genes (COG) identifiers in the mobilome COG category. Only COG IDs that are significantly enriched or depleted in pseudogenes vs intact genes across one of the ten tested species are shown (focused on the ultra-rare pangenome partition). These tests were run per-species and restricted to redundant elements. Log-odds ratios > 0 indicate that a COG ID is enriched in pseudogenes vs. intact genes. The boxplot features are defined as follows: the centre line represents the median; the lower and upper hinges of the boxplots correspond to the 25th and 75th percentiles; the lower and upper boxplot whiskers extend to the lowest and highest points, respectively, to a limit of 1.5 multiplied by the interquartile range from the closest hinge.

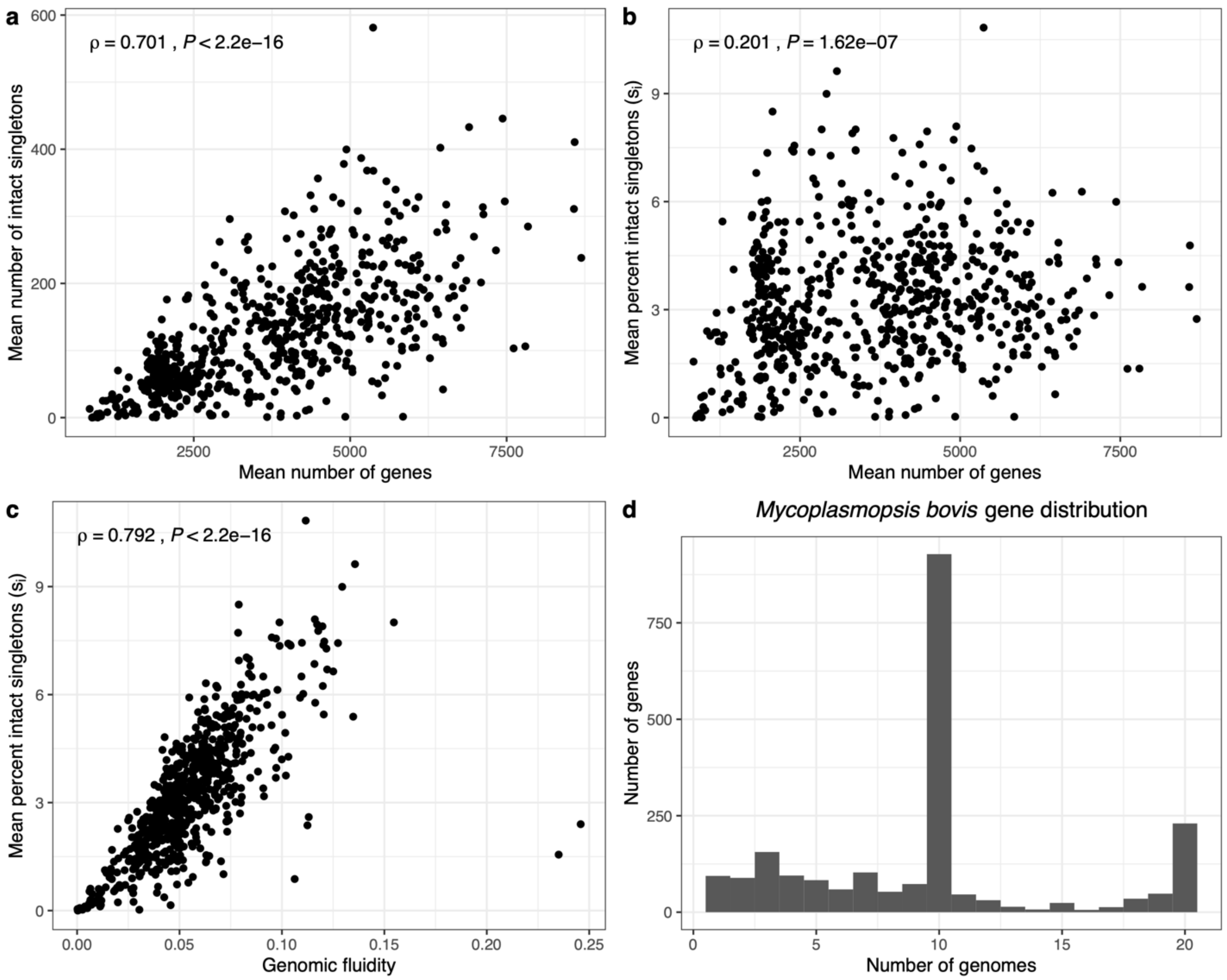


**Extended Data Figure 6**: Association of pangenome diversity metrics across 668 prokaryotic species. Panels a-c: Associations among different pangenome diversity metrics, where each point corresponds to one of the 668 species. Singleton-based metrics were estimated based on repeated subsampling to nine genomes per species. Spearman correlation coefficients and *P*-values are indicated. (d) Gene frequency distribution for 2,187 genes encoded by *Mycoplasmopsis bovis* genomes. This species is highlighted as it exhibited the highest genomic fluidity (right-most point on panel c, which is driven by population substructure (i.e. most genes are present at intermediate frequency, in 10/20 genomes).

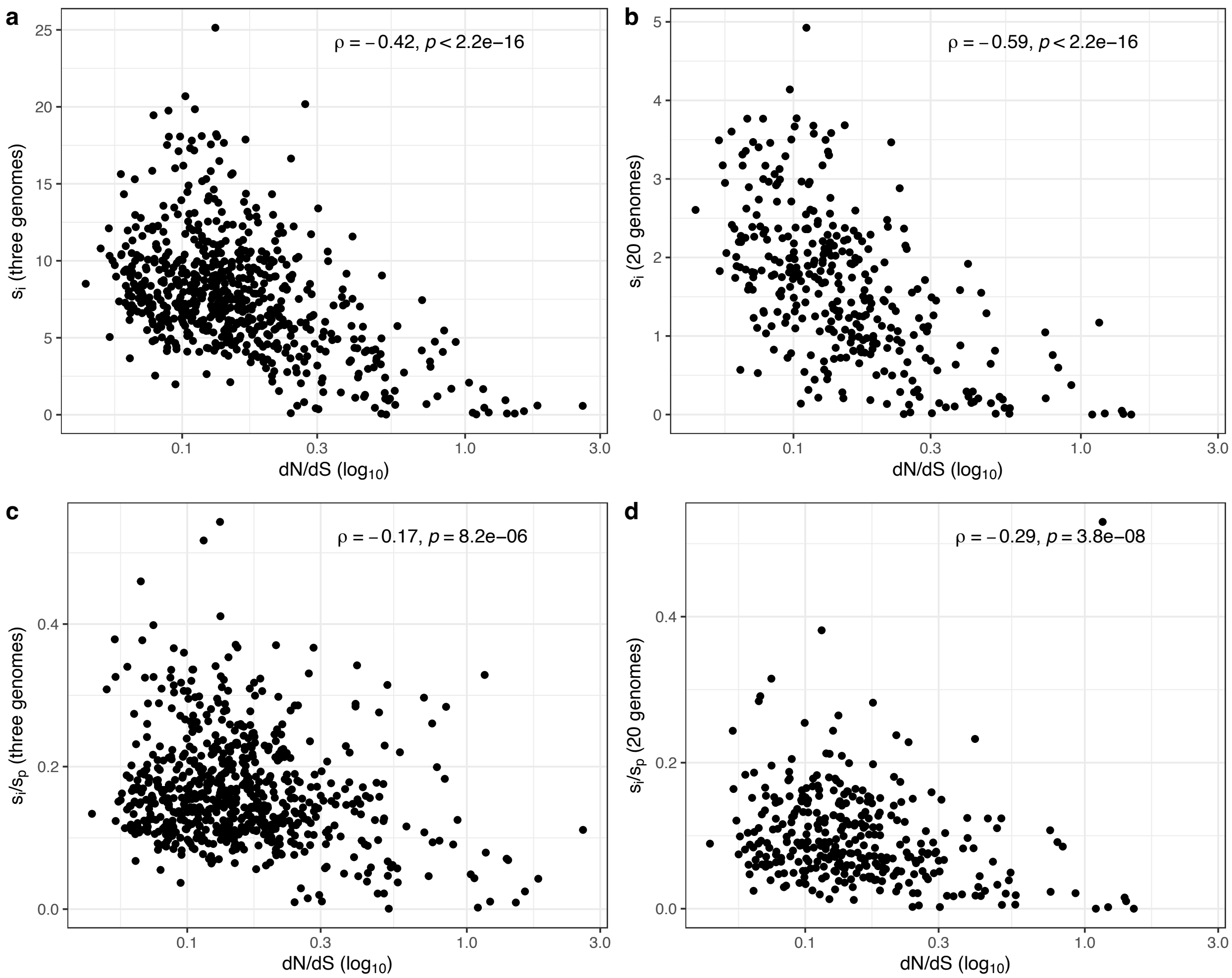


**Extended Data Figure 7**: Associations between dN/dS and pangenome diversity, as represented by repeated subsampling to three genomes to compute (a) $s_i$ and (c) and $s_i/s_p$, and repeated subsampling to 20 genomes to compute (b) $s_i$ and (d) and $s_i/s_p$. In the main text, $s_i/s_p$ is based on subsampling to nine genomes. Each point is one of 668 prokaryotic species. The Spearman correlation coefficients and *P*-values are indicated.

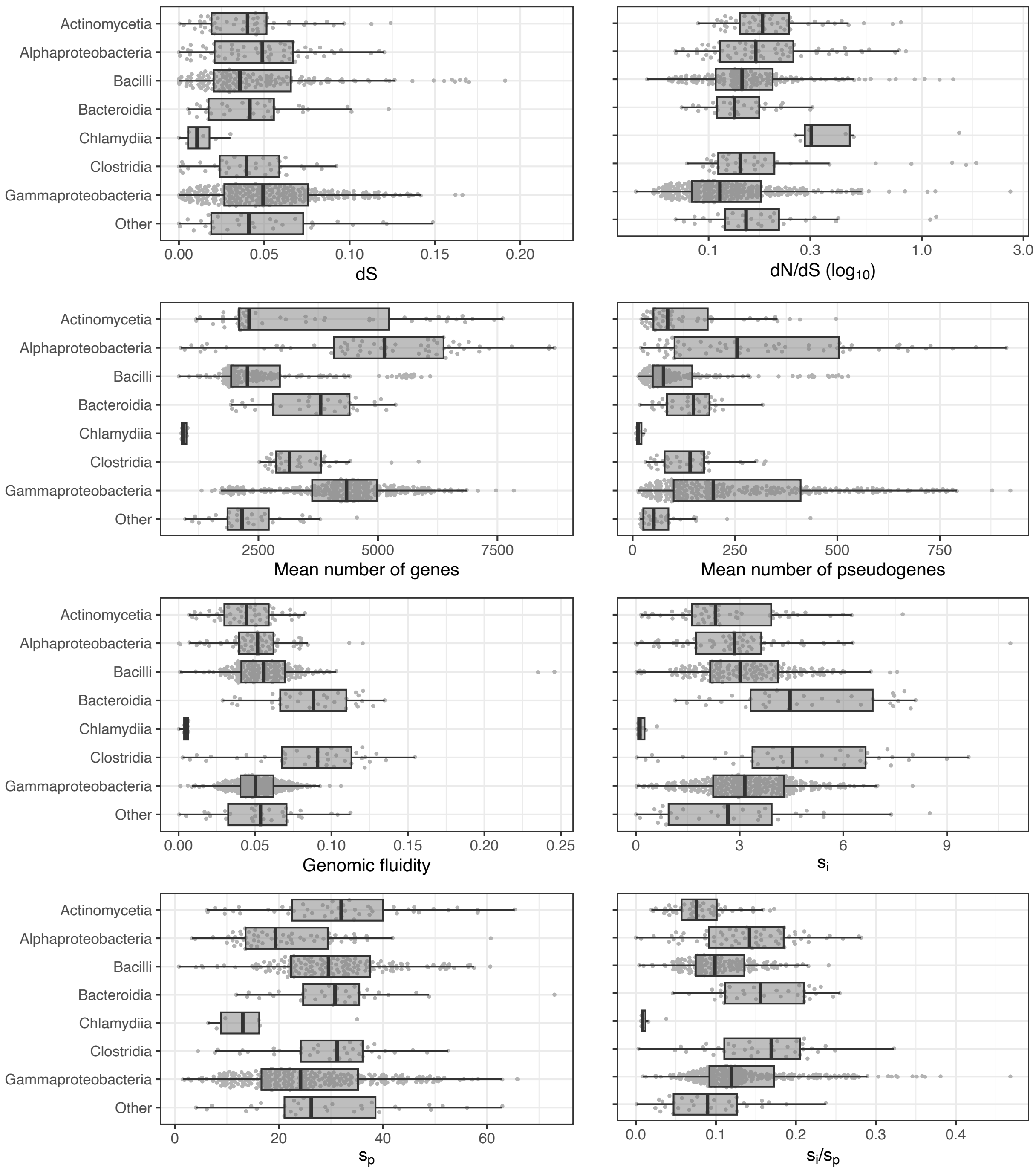


**Extended Data Figure 8**: Distributions of pangenome diversity and molecular evolution metrics stratified by taxonomic class (with classes with <= five species collapsed into 'Other'). Each point is a separate species.

| | | Mean no. genes | Genomic fluidity | $s_i$ | $s_i/s_p$ |
|---|---|---|---|---|---|
| Intercept coef. | Intercept | −0.872 | −0.074 | −0.231 | −0.530 |
| | Gammaproteobacteria | 1.225 | −0.136 | 0.108 | 0.692 |
| | Bacilli | 0.266 | 0.219 | 0.261 | 0.285 |
| | Alphaproteobacteria | 1.708 | −0.002 | 0.188 | 0.832 |
| | Actinomycetia | 0.767 | −0.179 | 0.127 | −0.039 |
| | Clostridia | 0.668 | 1.071 | 1.058 | 1.019 |
| | Bacteroidia | 0.835 | 1.146 | 1.038 | 1.089 |
| | Chlamydiia | −0.960 | −2.142 | −1.460 | −1.778 |
| Slope coef. | dS | −0.091 | 0.368 | 0.256 | −0.049 |
| | dN/dS | −0.026 | −0.137 | −0.359 | −0.196 |
| Model fit | Adjusted $R^2$ | 0.269 | 0.404 | 0.420 | 0.197 |

P−value
< 0.05
>= 0.05

No. species in class
300
200
100
0

**Extended Data Figure 9**: Summaries of four pangenome diversity linear models. One model was fit for each pangenome diversity metric: the mean number of genes, genomic fluidity, the percentage of singleton intact genes ($s_i$), and the ratio of the percentages of singleton intact genes vs. pseudogenes ($s_i/s_p$). All continuous response and predictor variables were standardized (i.e. converted to z-scores) prior to building models. Continuous variables were also transformed to normal distributions prior to this standardization (see Online Methods). Coefficients are displayed for each model, split by those that affect the intercept vs. the slope. The adjusted $R^2$ is also indicated for each model, and the cell colouring indicates whether each value is statistically significant ($P < 0.05$). The number of genomes per taxonomic class is indicated by the blue bar. The category used to infer the overall intercept was based on a combination of all classes with <= 5 species present. Note that Chlamydiia is the class, not the common genus *Chalmydia*. These models were built based on 667 species, after excluding one species with no singleton intact genes, and contained 657 degrees of freedom.

### Extended Data Table 1 – Summary of 10 species used for in-depth pangenome analysis

| Species | Total genomes | Total genes | Total pseudo. | Total intact clusters | Total pseudo. clusters | Mean intact clusters per genome | Mean pseudo. clusters per genome | Mean intact gene cov.[a] per genome | Mean pseudo. cov. per genome |
|---|---|---|---|---|---|---|---|---|---|
| *Agrobacterium tumefaciens* | 223 | 1076271 | 82318 | 24731 | 4464 | 4824.66 | 368.92 | 84.63 | 2.00 |
| *Enterococcus faecalis* | 1298 | 3339181 | 109584 | 26867 | 6257 | 2563.58 | 83.85 | 79.67 | 0.91 |
| *Escherichia coli* | 2955 | 12971835 | 2118815 | 79395 | 38196 | 4366.27 | 714.13 | 81.07 | 3.63 |
| *Lactococcus lactis* | 135 | 310580 | 8170 | 15485 | 1994 | 2286.21 | 60.17 | 80.39 | 0.68 |
| *Pseudomonas aeruginosa* | 4115 | 21908725 | 909029 | 95209 | 35636 | 5314.58 | 220.15 | 78.97 | 1.27 |
| *Sinorhizobium meliloti* | 166 | 1001437 | 80305 | 36736 | 9996 | 6002.48 | 483.39 | 80.58 | 2.14 |
| *Staphylococcus epidermidis* | 447 | 977677 | 33981 | 12681 | 2901 | 2181.84 | 75.05 | 77.97 | 0.87 |
| *Streptococcus pneumoniae* | 6845 | 11106685 | 1725479 | 37371 | 33803 | 1618.94 | 251.15 | 69.34 | 3.27 |
| *Wolbachia pipientis* | 716 | 642248 | 40310 | 4118 | 1336 | 895.28 | 55.45 | 73.09 | 1.29 |
| *Xanthomonas oryzae* | 326 | 1081258 | 128057 | 12426 | 6125 | 3177.30 | 387.67 | 72.00 | 2.94 |

[a]Percentage of total genome covered by specified elements.
